# Assessing the subarachnoid space anatomy on clinical imaging: utilizing normal and pathology to understand compartmentalization of the subarachnoid space

**DOI:** 10.1101/2024.05.05.592615

**Authors:** Khaled Almohaimede, Yusuf Alibrahim, Abu Bakar Butt, Pejman Maralani, Chris Heyn, Anish Kapadia

## Abstract

**BACKGROUND:** The goad of the study is to use CT imaging in patients with aSAH to evaluate the anatomic distribution of hemorrhage and compartmentalization of subarachnoid space to investigate potential in vivo visualization of recently discovered layer named subarachnoid lymphatic-like membrane (SLYM).

**METHODS:** We conducted a retrospective cohort study of cases with aneurysmal SAH (aSAH) at our institution between January 2015 and June 2022. Subarachnoid hemorrhage distribution into superficial and deep subarachnoid spaces was classified based on proximity to the dural or pial surfaces, respectively, as seen on multiplanar CT head.

**RESULTS:** A total of 97 patients with aSAH were included. Patients with lower modified Fisher score (MFS) of 1-2 were more likely to have SAH compartmentalizing in the “deep” pial-adjacent subarachnoid space. Patients with higher MFS of 3-4 were more likely to have SAH in both “superficial” and “deep” compartments along the brainstem. There is a significant association between the severity of aSAH - quantified by the MFS - and the distribution of the blood. Patients with higher MFS scores were roughly 7.6 times (p-value = 0.049) more likely to have hemorrhage at the “Superficial” juxta-dural subarachnoid compartment when compared to those with lower MFS scores.

**CONCLUSION:** This study suggests an imaging correlate to the recently discovered “SLYM”, potentially influencing aSAH compartmentalization, particularly in low-grade bleeds. While compartmentalization is limited in high-grade cases, these findings warrant further investigation with advanced imaging techniques to validate this membrane’s role and potential impact on CSF flow and aSAH pathophysiology.

## Introduction

The concept of subarachnoid space dates back to the 17th century, when Humphrey Ridley – a British physician-first described the basal cisterns such as the cerebellomedullary, quadrigeminal, and olfactory cisterns. He provided evidence for the presence of the arachnoid membrane as a separate meningeal layer and detailed how this membrane enveloped different cerebral vessels and intracranial nerves.^1,2^ Until recently it was accepted that the subarachnoid space represented a continuous space filled with cerebrospinal fluid (CSF)^3,4^. However, a recent publication authored by Mollgard et al has revealed the existence of a mesothelial layer that partitions the subarachnoid space into two distinct compartments.^5^ This study has provided evidence of this layer in both mice and human brains, and the authors have introduced the term “subarachnoid lymphatic-like membrane” (SLYM) to describe it. The SLYM acts as a semipermeable barrier, preventing the passage of molecules larger than 3 Kilodaltons, and contains myeloid cells, which play a role in innate immune function.^5^

It is currently believed that the “SLYM” envelops the brain^5^. However, the precise anatomical reflections of this membrane are not clearly understood. The SLYM is difficult to study histopathologically as the mesothelial layer is thought to fragment during dissection of cadaveric specimens. This is the reason the authors of the aforementioned study postulated why it remained undiscovered until recently. Given the fragility of the membrane, in vivo assessment of the anatomy using imaging may be an optimal way to target visualization. In vivo assessment with iodinated contrast or gadolinium-based contrast agents is not feasible due to their small molecular size^6,7^. Hence, leveraging of contrast provided by the pathological presence of subarachnoid blood in the setting of acute aneurysmal subarachnoid hemorrhage (aSAH)^8,9^ was postulated as the SLYM would be impermeable to the cellular contents of blood. As such, the current study aims to assess whether imaging can be used to evaluate the compartmentalization of the subarachnoid space and indirectly deduce the anatomic reflection of the SLYM utilizing acute aSAH as a model.

## Materials and Methods

### Patients’ selection

This retrospective single-center study was approved by the research ethics board with waiver of informed consent. All head CT scans with aSAH between January 2015 and June 31 2022 were identified by searching the departmental imaging database using specific keywords from CT reports (including “subarachnoid hemorrhage” and “aneurysm”). All adult (> 18-year-old) patients who presented with acute aSAH and had baseline routine plain head CT with multiplanar reformats were included. The cases with no sagittal reformats, poor visualization of the posterior fossa due to significant artifacts, or non-aneurysmal SAH (i.e. AVM, dAVF, Trauma) were excluded. Patient demographics, World Federation of Neurosurgical Societies grading system, aneurysm location, aneurysm size and type were retrieved using electronic medical records.

### Image acquisition and analysis

All the standard-of-care head CTs were performed on 64-section CT scanners (Siemens Healthcare, Forchheim, Germany). Unenhanced CT scans covered the area from the skull base to the vertex with multiple reformats.

The initial unenhanced CT head was used to grade the subarachnoid hemorrhage using the modified Fisher scale (MFS). Additionally, the midline sagittal images were used to classify the distribution of subarachnoid blood within the basal cisterns into predominantly “Superficial” or predominantly “Deep” subarachnoid space based on the proximity of the hemorrhage to the dural or pial surfaces, respectively. Further classification was done based on the location in the interpeduncular, pre-pontine, pre-medullary cisterns and peri-spinal CSF space at the foramen magnum. The subarachnoid space within the cerebral convexities and fissures were classified into prominent or narrow, based on the degree of volume loss, and further assessment for compartmentalization was done only if the space is prominent (i.e. those demonstrating volume loss). CTA scans were used to assess the location, size, and shape of the aneurysms.

### Statistical Analysis

Statistical analysis was performed using SPSS (version 26.0) IBM Corp. Released 2019. IBM SPSS Statistics for Windows, Version 26.0. Armonk, NY: IBM Corp. (n.d.). [Computer software]. and R (version 4.3.1). Descriptive statistics for patients’ demographic data were generated. The modified Fischer Scores were dichotomized into two groups; Low (Score 1-2), and High (Score 3-4). The relationship and distribution of SAH at different cisterns along the dural and pial surfaces with the degree of MFS was assessed using Chi-Square correlation test. Statistical significance was set at p-value < 0.05.

### Results

A total of 97 patients with aSAH were included with and average age of 60.36 +/-12.1 years (28-89) composed of 75 (77.3%) females and 22 (22.7%) males. The majority (72.2%) of subarachnoid hemorrhage cases had high MFS (3-4), with the remaining 27.8% having low MFS (1-2).

A total of 63 cases (64.9%) had the hemorrhage present in the “deep” juxta-pial subarachnoid space along the brainstem only, whereas no cases were seen only in the “superficial” juxta-dural subarachnoid space. Furthermore, 26 cases (26.8%) had hemorrhage in both the “superficial” and “deep” compartments along the brainstem. There were 8 cases (8.2%) with no subarachnoid hemorrhage along the brainstem but had hemorrhage along the cerebral convexities.

There is a significant association between the severity of aSAH - quantified by the MFS - and the distribution of the blood. Patients with higher MFS scores were roughly 7.6 times (p-value = .049) more likely to have hemorrhage at the “Superficial” juxta-dural subarachnoid compartment when compared to those with lower MFS scores.

At low MFS, hemorrhage present at the “Deep” juxta-pial subarachnoid space at any location along the brainstem is significantly associated with subarachnoid hemorrhage at other “Deep” juxta-pial locations (Table 1). For example, “Deep” juxta-pial aSAH at the pre-medullary cistern is significantly associated with juxta-pial blood at the prepontine (χ2 = 11.2; p-value = 0.001) and peri-spinal cisterns (χ2 = 5.3; p-value = 0.021). However, we found no cases at low MFS with “Superficially” distributed blood along the Juxta-dural subarachnoid space at the peri-spinal or pre-medullary cisterns and only 3 cases in the prepontine cistern but without statistically significant associations of aSAH with other locations (p value >0.077, 0.964, 0.233). On the other hand, high mFS aSAH was found to distribute along the “superficial” or “deep” compartments or both. For example, juxta-pial peri-spinal aSAH was significantly associated with juxta-pial pre-medullary (χ2 = 10.1; p-value = 0.001) and juxta-dural peri-spinal and pre-medullary (both χ2 = 4.8; p-value = 0.028) aSAH. Similarly, juxta-dural prepontine aSAH was significantly associated with juxta-dural premedullary (χ2 = 10.3; p-value = 0.001) and peri-spinal (χ2 = 5.0; p-value = .025) aSAH.

**Table 1.**
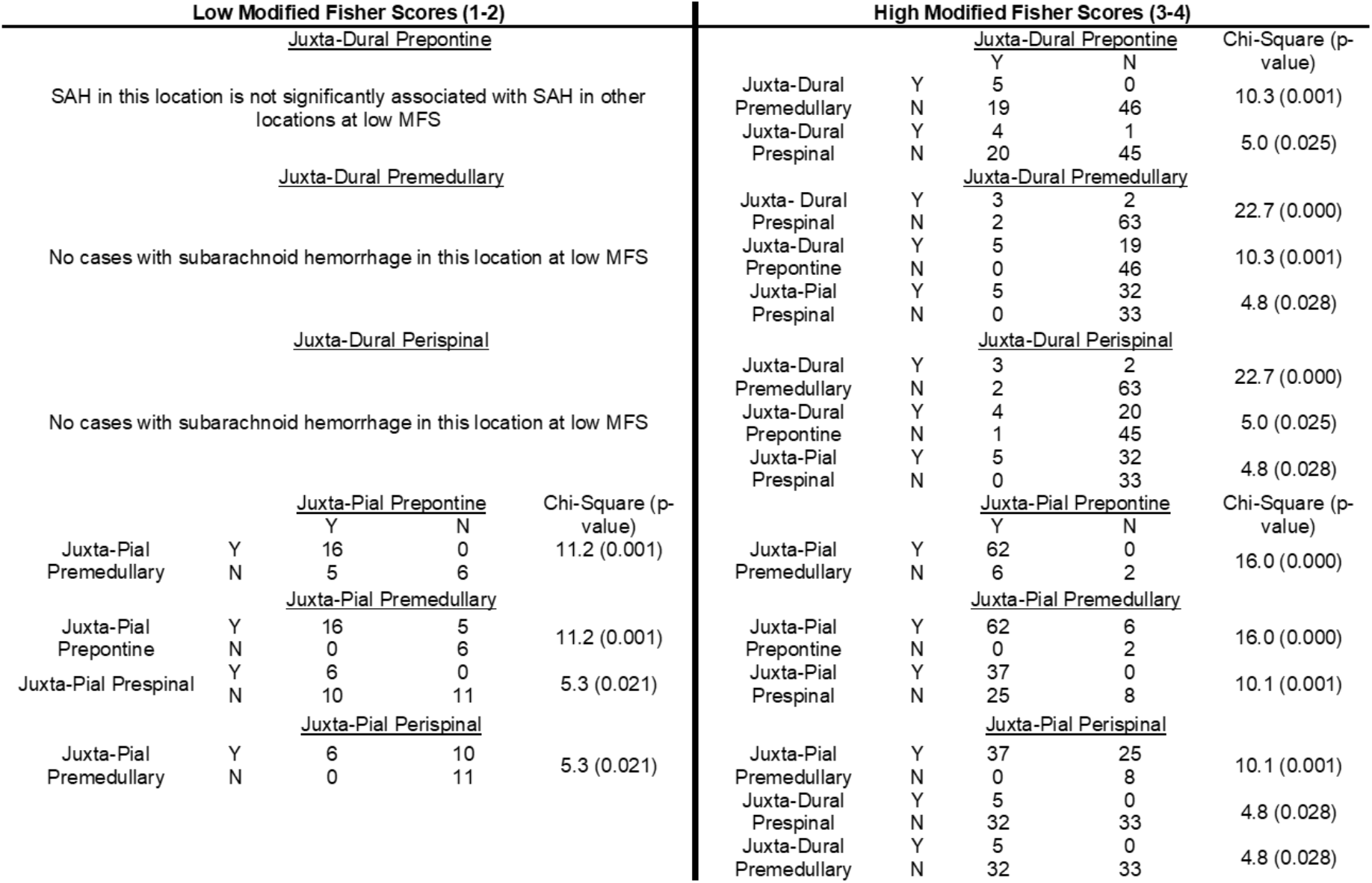
Relationship and associations of hemorrhage at different locations along the dural-adjacent and pial-adjacent subarachnoid.

## Discussion

The current study is the first in-vivo study to evaluate for the compartmentalization of the subarachnoid space, leveraging the distribution of blood products in the setting of acute aSAH. Our results suggest a potential imaging-based correlation to the recently discovered SLYM, which partitions the subarachnoid space into two distinct compartments. This correlation appears particularly evident in lower-grade aSAH (MFS 1-2), where blood functions as an internal contrast agent, highlighting the suspected location of this membrane. Conversely, in cases with more extensive hemorrhage (MFS 3-4), visualizing the partition layer becomes challenging. We hypothesize that this could be due to membrane displacement or rupture secondary to the high blood volume and elevated pressure within these confined spaces (Figure 1).

**Figure 1.**
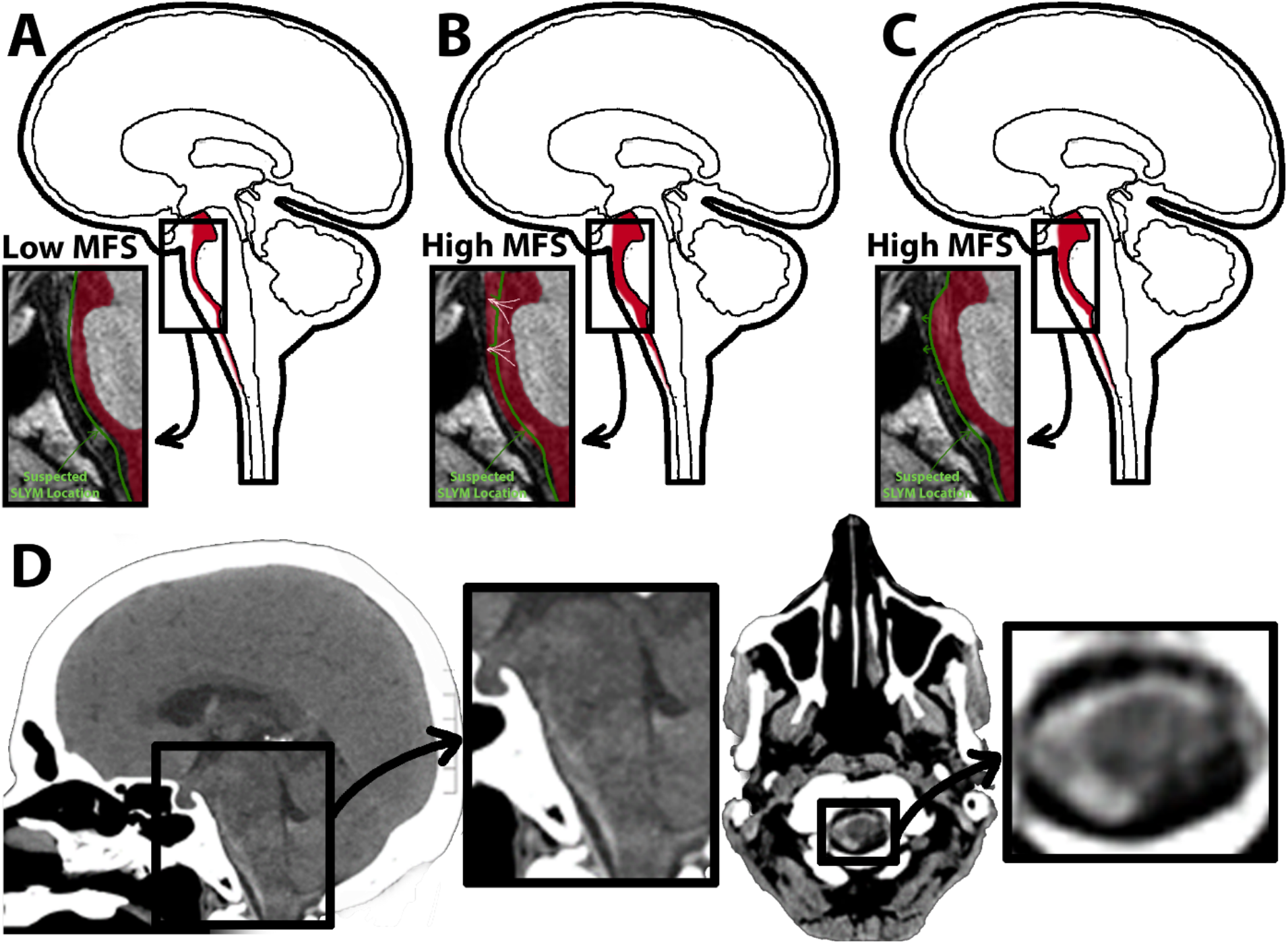
Illustration depicting the compartmentalization of aneurysmal subarachnoid hemorrhage in high and low modified Fisher scale (MFS). **(A)** Patients with low MFS of 1-2 are more likely to have SAH compartmentalize towards the deeper or pial-adjacent subarachnoid space, which is hypothesized to be related to the SLYM acting as a barrier. **(B)** Patients with higher MFS of 3-4 are more likely to have SAH abutting both pial and dural surfaces, which can be explained by the rupture of the SLYM at higher pressures allowing the subarachnoid blood deep to the SLYM to distribute superficial to the SLYM. **(C)** As the SLYM is not directly visualized in CT head, another explanation for the apparent redistribution of subarachnoid hemorrhage towards the dural surface is the displacement of the SLYM towards the dura. **(D)** Midline sagittal CT brain demonstrating SAH along the pial surface of the premedullary cistern. In the prepontine cistern blood abuts the pial and dural surfaces. Axial CT brain at the level of foramen magnum/upper cervical cord shows semi circumferential SAH along the pial surface of the cord.

Identifying this membrane and understanding its role could have significant implications for various neurological disorders. Elucidating its influence on CSF circulation could shed light on related pathologies. Future investigations utilizing advanced imaging techniques with higher resolution are warranted to further validate these observations and refine our understanding of this newly described anatomical structure. Such advancements could pave the way for improved diagnostic accuracy and potentially inform future therapeutic strategies related to aSAH and other neurological conditionas.

## Conclusion

This study suggests an imaging correlate to the recently discovered SLYM membrane, potentially influencing aSAH compartmentalization, particularly in low-grade hemorrhage. While compartmentalization is limited in high-grade cases, these findings warrant further investigation with advanced higher resolution imaging techniques to validate this membrane’s role and potential impact on CSF flow and aSAH pathophysiology.

